# Astrocyte-oligodendrocyte crosstalk dependent myelination via secreted protein YKL40

**DOI:** 10.64898/2026.09.08.750142

**Authors:** Oscar A. Domnguez, Soha Munir, Surya Chandra Rao Thumu, Connor Tuck, Jean Patrick Gonzales, Unsong Oh, Tomasz Kordula, Babett Fuss, Sandeep K. Singh

**Author notes:** Correspondence: Sandeep K. Singh, PhD.

## Abstract

Although oligodendrocyte differentiation and myelin formation are inherent properties of oligodendrocyte progenitor cells (OPCs) that are guided by intrinsic transcriptional and epigenetic programs, this process in the brain is finely regulated by signals coming from other cells, including astrocytes. Here, we identified the astrocyte secreted protein YKL40 at the center of astrocyte-OPC cross-communication and myelination in the developing brain. We find that YKL40 is expressed by astrocytes within white matter areas in the developing brain, coinciding with the OPC differentiation and myelination. Deletion of YKL40 in astrocytes showed delayed developmental myelination and reduced OPC proliferation. Interestingly, coculture with OPCs *in vitro* specifically induces YKL40 expression in astrocytes, which in turn promotes OPCs’ differentiation and OPC proliferation. Mechanistically, purified YKL40 significantly induced the expression of transcription factors Olig2 and MYRF (Myeline regulatory factor) in OPCs. Therefore, we identified a novel mechanism of OPC-astrocyte crosscommunication dependent myelination by astrocytic YKL40 in the developing brain.

## Introduction

The proper development of myelin sheaths is crucial for establishing functional and efficient neuronal circuits^1–3^. Alterations in myelin structure or function underlie the pathophysiology of many neurological disorders^1^. Myelin sheaths are primarily synthesized by oligodendrocytes (OL) in the central nervous system (CNS) and involve a tightly regulated and coordinated differentiation process of oligodendrocyte precursor cells (OPCs)^3,4^. In the rodent brain, OL differentiation primarily occurs postnatally between P4-P21 and comprises of three distinct phases: (1) the proliferation of NG2^+^/ chondroitin sulphate proteoglycan 4 (CSPG4^+^) and PDGFRα^+^ OPCs during P2-P5, (2) the conversion of OPCs to O4+/ low PDGFRα^+^ pre-oligodendrocytes (POL) during P5-P9, and (3) the differentiation of POLs to myelinating oligodendrocytes post-P10, which is marked by the expression of myelin basic protein (MBP), myelin regulatory factor (MYRF), and myelin oligodendrocyte glycoprotein (MOG)^3–5^. The timing of OL differentiation is critical for proper myelin formation in the developing CNS; therefore, this process is highly regulated by internal factors that are intrinsic to OL-lineage cells and extracellular signals from other cells. Others have shown that several transcriptional regulators, such as ID2, ID4, Nkx2.2, Olig1, Olig2, Sox10, and MYRF, act as primary intrinsic cues to modulate OL differentiation^1,2,4,6^. Furthermore, extracellular cues, such as Notch, BMP, Shh and Wnt/β-catenin act as negative regulators of myelination, while LIF, CNTF and BDNF signaling enhance OL differentiation^7^. Surprisingly, the precise mechanisms of how extracellular signals and intrinsic factors cooperate during myelination and development has not been fully elucidated.

Interestingly, OPCs are generated by the same neural progenitor cells that give rise to neurons and astrocytes in the CNS^2,3^. OL differentiation and myelination processes coincide with the differentiation and maturation phase of astrocytes, as well as neuronal arborization/circuit formation, implying cellular crosstalk between these cells to coordinate efficient myelination. Recently, neural activity has emerged as a key phenomenon regulating OL differentiation and myelination^8–10^. Early in the developing rodent brain, OPCs migrate along the vasculature to disperse into their destinations, and detachment from endothelial cells permits their subsequent differentiation^11,12^. Interestingly, astrocyte end foot formation aids in OPC detachment from the endothelium, enabling their differentiation^12^. Similarly, astrocytes induce OL proliferation and differentiation by secreting the trophic factors, PDGF, LIF, CNTF, BDNF and BCAN^13–17^. In the optic nerve CNTF expression in astrocytes starts at the onset of myelination to facilitate the final differentiation of OPCs^16,18,19^. Furthermore, local ablation of astrocytes in the developing spinal cord dramatically reduced mature OLs and inhibited myelination^20^. Although much is known about the role of astrocytes during myelination, there is a deficit in understanding the dynamic interactions between astrocytes and OPCs to fine-tune myelination. For example, does OPC- astrocyte crosstalk regulate developmental myelination?

Here we report YKL40 (gene name *Chil1*) as a central player in OPC-astrocyte crosstalk-dependent myelination. YKL40, also known as CHI3L1 (Chitinase-3- Like1), is a secreted glycoprotein that belongs to 18-glycosyl-hydrolase family of proteins but without the glycosyl hydrolase enzymatic activity due to mutations in the active site^21,22^. YKL40 is expressed by many cell types including macrophages, chondrocytes, neutrophils and synovial cells outside the CNS, and astrocytes within the CNS. It can elicit a multitude of actions including cell proliferation, immune cell activation, extracellular matrix remodeling and cell migration. It exerts many of these functions via interacting with several receptors, including IL13Rα2, CD44, and RAGE^22^. Moreover, increased expression of YKL40 highly correlates with inflammation and cancer^22^. Elevated levels of YKL40 in cerebrospinal fluid have been associated with aging and is considered as one of the biomarkers in Alzheimer’s Disease (AD) and multiple sclerosis (MS)^23,24^. Moreover, global YKL40 or astrocyte-specific YKL40 knockout mice are protected against plaque burden and exhibit restored memory functions^25,26^. We have also previously shown that YKL40 expression increases during astrocyte differentiation and in astrocytes exposed to proinflammatory cytokines^27,28^.

Although YKL40 is upregulated in many neuropathologies and has emerged as a novel therapeutic target, its precise role in the developing and diseased brain is not known. Here we examined the expression and function of YKL40 in the developing brain. Surprisingly we discovered that OPC-astrocyte coculture leads to enhanced expression of YKL40 in astrocytes and subsequent stimulation of OPC differentiation. Further, we generated astrocyte specific YKL40 knock out mice which presented delayed developmental myelination. Interestingly, YKL40 affects developmental myelination by acting at two distinct stages: 1) by regulating OPC proliferation and 2) by modulating oligodendrocyte maturation. Our findings establish YKL40 as a central player in OPC-astrocyte crosstalk-dependent myelination.

## Materials and Methods

### Mice

All mice used in this study were maintained on the C57BL/6J background. Knockout mice were generated by crossing YKL40 loxP/loxP mice (#T013652, GemPharmatech) with *GFAP-cre* mice (The Jackson Laboratory, stock# 024098). Mice were housed in cages, 2-5 mice per cage, in temperature and humidity-controlled conditions. Male and female mice were collected for this study. The mice used for this study are WT C57/BL6, ALDH1L1-GFP, and YKL-40 f/f GFAP-Cre (YKL40).

### Transcardial Perfusion and Cryopreservation

Mice were sedated with isoflurane and perfused transcardially with 1xTBS pH7.4. Brains used for RNA and protein isolation were flash frozen in liquid nitrogen and stored at -80 °C. Brains used for immunohistochemistry were fixed by perfusing with 4% PFA, left in 4% PFA overnight, washed 3 times in 1xTBS, then left in a 30% sucrose solution made in 1xTBS for 2 days. Fixed brains were then cryopreserved on dry ice in a 1:1 OCT:30% sucrose solution, then stored at -80°C. Cryopreserved brains were cut into sagittal and or coronal sections of 15-20 µm on a cryostat and brain slices were placed on slides and stored at -80°C until use.

### Tissue Immunofluorescence

Brain sections were quickly washed in 0.2% TBS-T (1x TBS with 0.2% Triton X-100). Sections were then incubated in blocking buffer (10% NGS made in 0.2% TBST) for 1 hour at room temperature. Sections were then incubated in primary antibody solutions made in blocking buffer, then stored in a humidified chamber at 4 °C overnight. Primary antibodies used were chicken GFAP (AB2313547, Aves Labs, 1:1000), chicken Olig2 (MABN50 Olig2, EMD Millipore, 1:500), guinea pig Aquaporin 4(429 004 AQP4, Synaptic Systems, 1:500), guinea pig GLT1 (AB1783 GLT1, EMD Millipore Corp, 1:1000), rabbit Iba1 (019-19741 Iba1, Fujifilm, 1:500), rabbit NeuN (#12943, Cell Signaling Tech, 1:500), rat MBP (MAB5272 MBP, EMD Millipore Corp, 1:50), rabbit NG2(1:200), mouse CC1 (MABC200, EMD Millipore, 1:500), Ki67 (#9129, Cell Signaling Technology, 1:500), mouse CNPase (C5922, SIGMA, 1:500). Sections were then washed in 0.2% TBS-T 3 times for 10 minutes each. They were then incubated in secondary antibody solution made in blocking buffer for 2 hours, all done in dim light conditions. Secondary antibodies used are anti-rabbit 594 (A32740 Anti-Rabbit 594, Invitrogen), anti-chicken 488 (A11039 Anti-Chicken 488, Invitrogen), anti-mouse 488 (A32732 Anti-Mouse 488, Invitrogen), and anti-rat 647 (A21247 Anti-Rat 647, Invitrogen), all made in a 1:500 solution. Sections were then washed in 0.2% TBS-T 3 times for 10 minutes each, done in dim light. Sections were sealed in VECTASHIELD (Vector labs) mounting media with DAPI. Slides were imaged using a Zeiss LSM 710 or LSM 880 confocal microscope.

### RNA Isolation and qPCR

Brain tissues were collected and flash frozen as described previously and were mechanically crushed into powder on dry ice using a pulverizer. A portion of the tissue was lysed with Trizol following manufacturer’s protocol. Human astrocytes and mouse OPCs cultures were lysed with Trizol following manufacturer’s protocol. RNA was isolated using the Nucleospin RNA Isolation Kit (Machery Nagel, REF 740955). RNA was reverse transcribed using the Promega RQ1 DNAse protocol and the High-Capacity cDNA Reverse Transcription Kit (4368813, Thermo Fisher). Samples were run in duplicates and normalized against GAPDH gene expression and fold change was quantified using ΔΔCt method. Primers used were from IDT (mouse Chil1, mouse Chia, mouse Chid1, mouse Chit1, mouse Chi3l3, mouse OVGP1, mouse GFAP, mouse CSPG4. mouse SMOC1, mouse ENPP6, mouse Dusp15, mouse MYRF, mouse MBP, mouse Olig2, mouse MOG, mouse GAPDH, human YKL-40, human GFAP, human GLT-1, human GLAST, and human GAPDH).

### Western Blotting

A portion of the frozen brain powder was lysed in a modified RIPA buffer (50 mM Tris pH 7.6, 150 mM NaCl, 1mM EDTA, 0.1% SDS, 1% sodium deoxycholate, 1% Triton X-100, 1:100 protease inhibitor). Protein concentration was estimated using a BCA assay and equal amount of protein lysate per samples was loaded in a gradient 4-15% acrylamide gel (BioRad). Gels were placed in an Invitrogen PowerBlotter Station to transfer the proteins to a nitrocellulose membrane. Membranes were washed in TBS-T (1xTBS with 0.1% Tween20) for 1 minute and blocked for 1 hr in 10% milk made in TBS-T. Membranes were incubated overnight with primary antibody solution made in 2% BSA in TBS-T. The primary antibodies used were goat YKL-40 (SC30465 GP-39, Santa Cruz Biotechnology, 1:1000), rat MBP (MAB5272 MBP, EMD Millipore Corp), mouse CNPase (C5922 CNPase, SIGMA, 1:750), mouse MOG (MAB5680, EMD Millipore Corp, :2000) and rabbit tubulin (2128S B-Tubulin (9F3), Cell Signaling Tech, 1:500). Membranes were washed three times in TBS-T before being incubated in a secondary antibody solution in 5% milk in TBS-T for 1 hour. The membrane was then washed three times in TBS-T to remove the milk. Membranes were visualized in an Azure 600 after incubating in a 1:1 luminol solution for 60 seconds and analyzed using ImageJ.

### Oligodendrocyte progenitor cell (OPC) isolation

OPCs were isolated using Milteny beads based on a protocol described in^29^ and as suggested by Milteny biotech. Cortices from P7 C57/BL6 WT pups were collected and dissected, digested in papain buffer (Papain, Worthington, LK003176) with DNAse, at 34°C water bath for 45 minutes. The papain activity was inhibited by Neurobasal media+10%FBS+DNAse and digested tissue chinks were triturated and passed through a 70-um filter and then spun down at 200 g for 11 minutes. The supernatant was aspirated, and the pellet was resuspended in Neurobasal media+10% FBS+DNAse and passed through a 40- µm filter and spun down at 200 g for 11 minutes. The cell pellet was then resuspended in ice cold MCS solution (1xPBS, 0.5% BSA, and 2 mM EDTA). The supernatant was aspirated, and the tissue pellet was resuspended in ice cold MCS, then incubated with an anti-Fc receptor antibody for 10 minutes. After this, it is incubated with anti-O4 magnetic beads (Milteny Biotech) at 4^0^C for 15 minutes which was then spun down at 200 g for 10 minutes. The supernatant was aspirated, and the cell pellet was resuspended in cold MCS. The magnetic separator column was attached to a magnetic stand. A 20-μm filter was placed on top of the column. The column was placed over a 50 mL falcon tube. The filter and column were washed with cold MCS buffer. The cell suspension and beads were passed through the 20-um filter and into the column. The filter was washed with MCS buffer. The filter was discarded, and the column was washed 3 times with MCS buffer and 1 time with OL proliferation media (Neurobasal media, 1% Pen Strep, 1% Glutamine, 1% Sodium Pyruvate, 1xB27, 20 ng/mL bFGF, 20 ng/mL PDGFaa). The column was taken off the magnetic stand and placed into a 15 mL falcon tube. OL proliferation media was added to the column, which is then pushed through the column using the plunger with the column. Cells were counted and then plated.

### Application of recombinant YKL-40 on Isolated OPCs

OPCs were isolated using the previously written protocol. Approximately 20,000 cells were plated onto a PDL and laminin coated 8 well chamber slide and approximately 100,000 cells were plated on a PDL and laminin coated 24 well plate. After 1 day, media was replaced with OL Differentiation media (Neurobasal media, 1% Pen Strep, 1% Glutamine, 1% Sodium Pyruvate, 1xB27, 40ng/ml T3). Varying concentrations of recombinant YKL-40 (0-200 ng/mL) were added for a total of two days. After another 24 hours, the media was replaced with new OL Differentiation Media with new YKL-40. At the end of 48 hours, cells were fixed with 4% PFA for immunofluorescence staining or Trizol lyzed for RNA isolation and gene expression analyses.

### Coculture of Human Astrocytes with Mouse OPCs

Human astrocytes were grown onto 10 cm^2^ culture treated plates in a 20% FBS containing complete media as described^30^. Two days after the media was removed and replaced by 10% FBS complete media. Between DIV4-6 human astrocytes were lifted off by trypsinization and resuspended in OPC differentiation medium and added at ∼100,000 per well on to the DIV1/2 OPC cultures in a 24-well plate. Three groups were made: Group 1, which is the astrocyte only control; group 2, which is mouse OPC only control; and group 3, which is the astrocyte-OPC cocultures. Two days later cells were lysed in Trizol and collected for RNA isolation, as described above.

### Cell Immunocytochemistry

OPCs were fixed and stained following the protocol described in Ippolito and Eroglu, 2010. Primary antibodies used were chicken Olig2 (MABN50 Olig2, EMD Millipore, 1:500), rat MBP (MAB5272 MBP, EMD Millipore Corp 1:50), rabbit NG2 (Sigma, 1:200), Ki67 (9129S, Cell Signaling Technology, 1:500), and mouse O4 (1:10, O4 hybridoma). Cells were sealed in VECTASHIELD mounting media with DAPI. Slides were imaged using Zeiss 710 confocal microscope.

### Statistics

Statistical analyses were done using GraphPad Prism. The significance between 2 groups was determined using an unpaired student t-test. The significance between more than 2 groups was determined using one way ANOVA. Significance was determined by having a P value < 0.05.

## Results

### YKL40 expression is developmentally regulated in the mouse brain

Although expression of YKL40 is upregulated in inflammatory conditions and during several neuropathological states^19,24^, not much is known about its expression and functions in the developing brain. Previously, using entire mouse brain homogenates, we showed that YKL40 expression gradually increases during postnatal developing brain between P2 and P21^27^. Because YKL40 belongs to a large family of proteins with glycosyl hydrolase active site, here we examined the expression of YKL40 and its several family members in the developing mouse forebrains and cerebella during postnatal days 2-25 (P2-P25) by quantitative PCR (qPCR) when astrocyte differentiation occurs concomitantly with oligodendrocyte development and synaptogenesis. While expression of YKL40 gradually increased between P2 and P25 in the cortex (Cx), homologues of YKL40 remained either unchanged (Chi1, Chi3l3) or are downregulated (Chid1, Chit1 and OVGP1) (Fig. 1A). As expected, we found a gradual yet robust increase in MBP expression between P2-P25 (Fig. 1B). Western blot analyses confirmed increased expression of YKL40, and myelin marker MBP in the forebrain (Fig. 1C). Interestingly, similar results were also observed in the cerebellar tissue (Fig. 1D-F). Taken together these results suggest that YKL40 expression coincides with developmental myelination and that it may regulate critical functions in developing brain.

**Fig 1.**
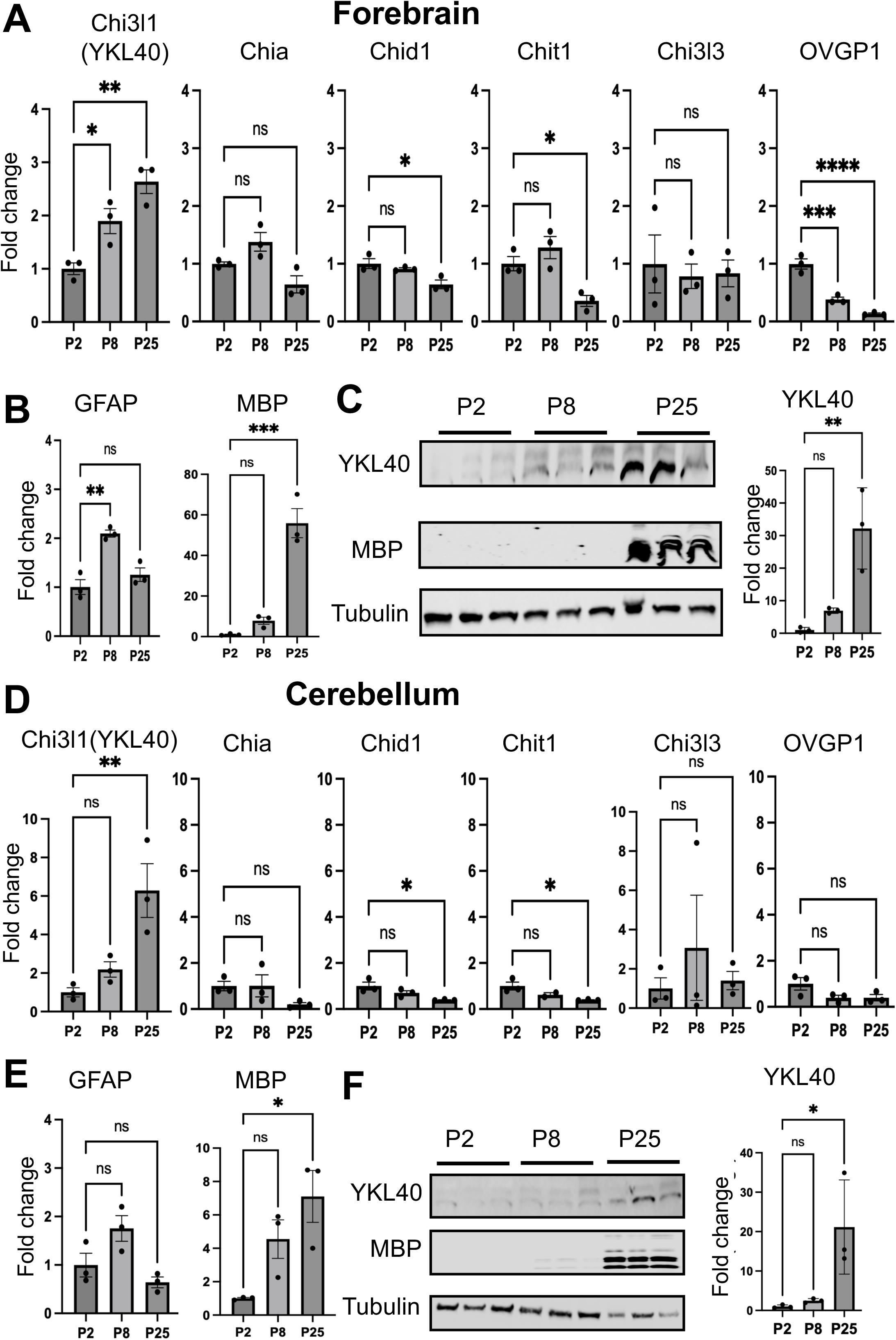
Dynamic expression of YKL40 family members in the developing mouse brain. Expression of YKL40 and family members in forebrains (**A-C**) and cerebellum (**D-F**). Quantitative real time PCR analyses of YKL40 family members (**A, D**) and markers of astrocyte (GFAP) and myeline (MBP) (**B, E**) at indicated postnatal age. Western blotting images and quantification of YKL40 and MBP (**C, F**) from the developing brain lysates. Tubulin is used as loading control. n=3 mice each group. Data mean ± SD, * P < 0.05, one way ANOVA.

### YKL40 is expressed specifically by maturing astrocytes in the mammalian brain

Previously we have shown that YKL40 is upregulated in maturing human astrocytes in a neural progenitor cell to astrocyte differentiation model^31^. To further explore the expression of YKL40 in the mouse brain, we immunostained YKL40 together with an astrocyte marker GFAP. Interestingly, a majority of cellular YKL40 staining colocalized with GFAP, confirming astrocytes as the major source of YKL40 in the brain (Fig. 2A). Intriguingly, we noticed that YKL40 signal is primarily localized within cortical, hippocampal and cerebellar white matter. On the other hand, grey matter (cortical) astrocytes lacked YKL40 signal (Fig. 2A). Since white matter is the primary region of oligodendrogenesis and myelination in the developing brain, we co-stained brain sections with YKL40 and oligodendrocyte specific markers including, Olig2 and CNP. As expected, Olig2 and CNP signal was enriched in the white matter areas and their expression was developmentally regulated (Fig.2B, C). While Olig2 positive staining decreased from P8 to P30, CNP staining increased during this time as reported by others^31^. More importantly, YKL40 signal intensity was also enriched in the white matter at P8 and P30. Remarkably, YKL40 positive cells were found around the Olig2/CNP positive cells, particularly at early developmental timepoints, P8, suggesting a possible role for OPCs in inducing expression of YKL40 in astrocytes. Moreover, we compiled data from various transcriptomic studies from human as well as rodent brains for the expression of YKL40 (Fig. S1A). Interestingly, YKL40 mRNA was among the top 20 genes expressed in mature human astrocytes compared to fetal human astrocyte^32^ corroborating our previous findings^27^. Similarly, YKL40 mRNA is most expressed by mature astrocytes in the rodent brains^33^. Significantly, YKL40 mRNA is being actively translated and highly enriched in astrocytes as measured by mRNA TRAP (translating mRNA affinity purification) approach (Fig. S1A bottom panel)^34,35^. Taken together, this data suggests that astrocytes are the primary source of YKL40 expression in the developing mammalian brain including rodents and human. Next, we performed RNA in situ hybridization (RNA FISH) to detect YKL40 mRNA in ALDH1L1-eGFP mouse brains where astrocytes are labeled by GFP. Indeed, we found YKL40 RNA signal localized within GFP expressing astrocytes in the corpus callosum (CC) and hippocampus (Fig. S1B and data not shown) further confirming YKL40 is mainly expressed by astrocytes in the mammalian brain.

**Fig 2.**
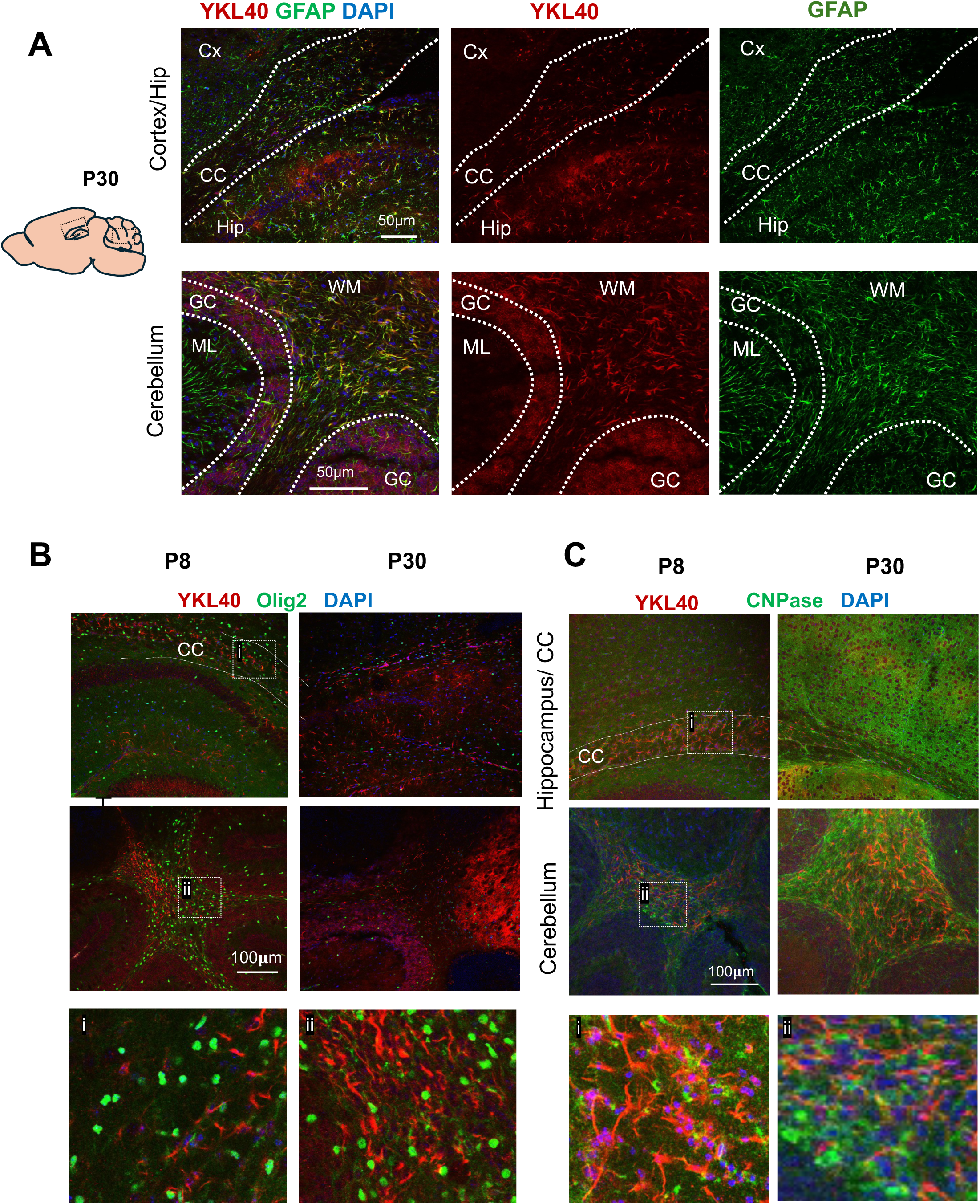
White matter astrocytes express YKL40 in the developing brain. YKL40 (red) and astrocyte marker GFAP (green) (**A**), or oligodendrocyte markers Olig2 (green) (**B**) and CNP (green)(**C)** in the mouse brain at P30 (**A**), or P8, and P30 (**B,C**). YKL40 colocalizes with the GFAP positive astrocytes within Olig2 and CNPase enriched brain areas including, corpus callosum (CC), cerebellum and hippocampus n=6 sections from 3 brain per time point.

### YKL40^ΔAST^ mice display delayed developmental myelination

Although YKL40 is upregulated in neuropathological conditions and affects neuroinflammation, its fundamental role in brain function during development and disease is not known. To address this, we generated astrocyte specific YKL40 knock out mice (YKL40^ΔAST^) by crossing YKL40 floxed mice (YKL40 fl/fl) with mice expressing Cre recombinase under mouse GFAP promoter (mGFAP-Cre line 77.6) (Fig. 3A). Quantitative real-time PCR showed ∼90% reduction in YKL40 mRNA in forebrains of YKL40^ΔAST^ compared to their WT littermates (Fig. 3B). Moreover, western blot analyses of YKL40 from astrocyte enriched cultures (with ∼90% astrocyte purity) of YKL40^ΔAST^ or littermate WT pups further confirmed decreased YKL40 expression (Fig. 3C). Similarly, YKL40 together with GFAP IF also confirmed YKL40 loss in astrocytes in this knock out model (Fig. 2D). Taken together this data establishes that YKL40 is specifically expressed by astrocytes in the brains at P30 mice. Because YKL40 is abundantly and specifically expressed by maturing astrocytes (Fig. 1, 2), we first asked if YKL40 is crucial for astrocyte development or functions. We stained P30 YKL40^ΔAST^ or littermate WT brains for major astrocyte marker proteins including GFAP and AQP4 and analyzed their expression in deep cortical layers (Fig. 2E, F). Overall intensity and percent area coverage of GFAP or AQP4 remained unaltered, suggesting no apparent defect in astrocyte development and/or function (Fig. 3E, F). Because YKL40 is a secreted protein and may interact with other CNS cells to exert its functions, we analyzed major neuronal, microglia and oligodendroglia markers including NeuN, Iba1 and MBP. We observed no obvious changes in NeuN and Iba1 marked cells (Fig. 3G-J). During these analyses, we noticed a noticeable reduction in the MBP expression (a marker of mature oligodendrocyte and myelin membrane) in YKL40^ΔAST^ brains at P30 (Fig. 4A). Hence, we next focused on characterizing the development of oligodendrocytes in YKL40^ΔAST^.

**Fig 3.**
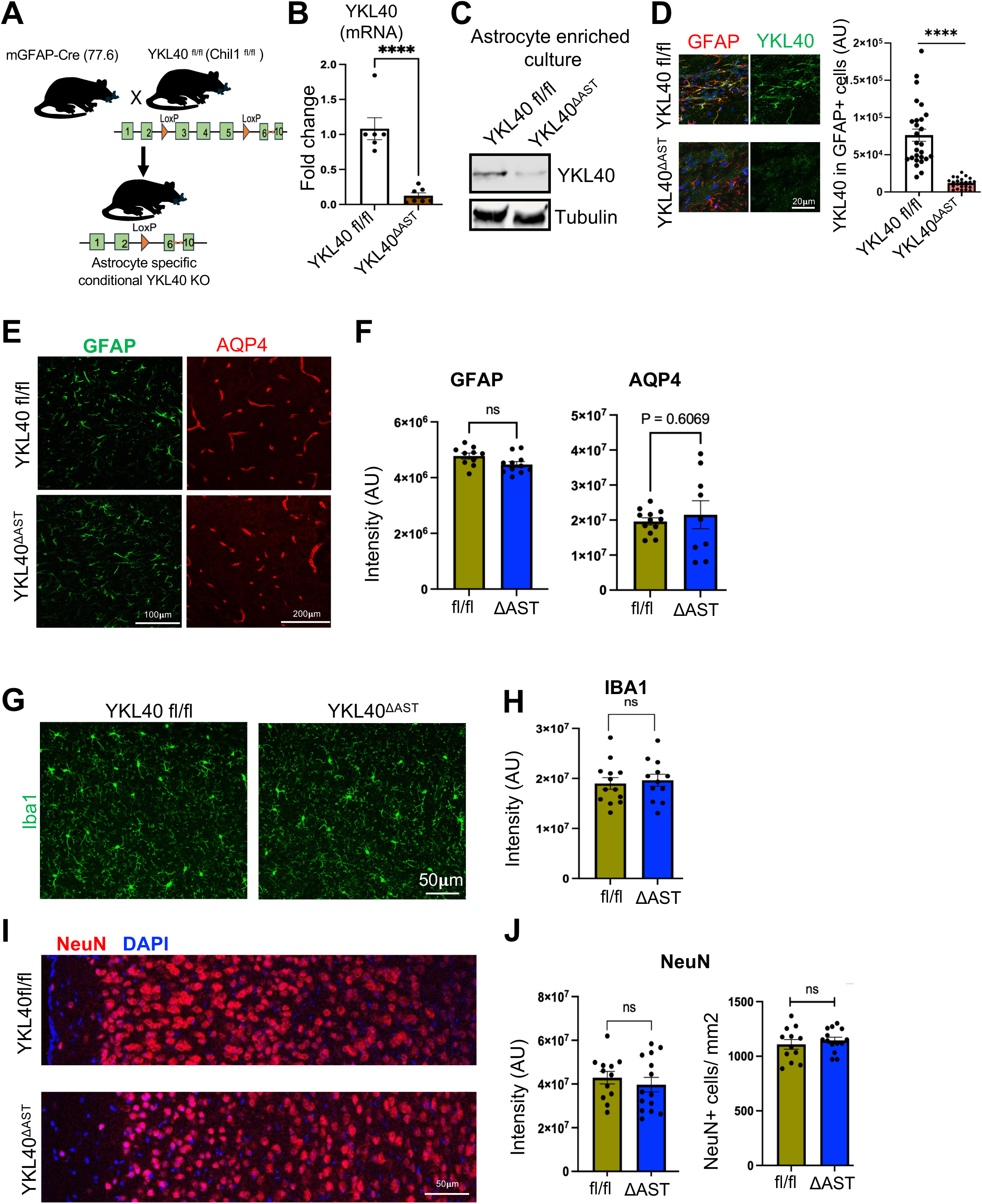
Generation and validation of astrocyte specific YKL40 knock out mice. (**A**) Scheme of astrocyte specific YKL40 knock out mice (YKL40^ΔAST^) by crossing YKL40 ^fl/fl^ (Chil1^fl/fl^) with mGFAP-Cre (77.6) line. (**B**) qPCR analyses of YKL40 from the YKL40^ΔAST^ or littermate control at P30 normalized to GAPDH. n=6, 7 mice per group (**C**) Western blot of YKL40 from astrocyte enriched cultures of YKL40^ΔAST^ or littermate control mice. (**D**) Confocal images and quantification of YKL40 (green) and GFAP (red) in the corpus callosum of P30 YKL40^ΔAST^ or littermate control mice. n= 20-20 astrocytes from 3 mice each. Confocal images (**E**) and their quantification (**F**) of mature astrocyte markers GFAP (green) and Aqp4 (red) from P30 YKL40^ΔAST^ or littermate control mice. Confocal images (**G, I**) and their quantification (**H,J**) of microglia marker Iba1 (green) (**H**) and neuronal marker NeuN (Red) (**I**) from P30 YKL40^ΔAST^ or littermate control mice. n=12-14 images from 3, 4 mice per group. Data mean ± SEM, * P < 0.05, unpaired t-test; ns=non-significant.

**Fig 4.**
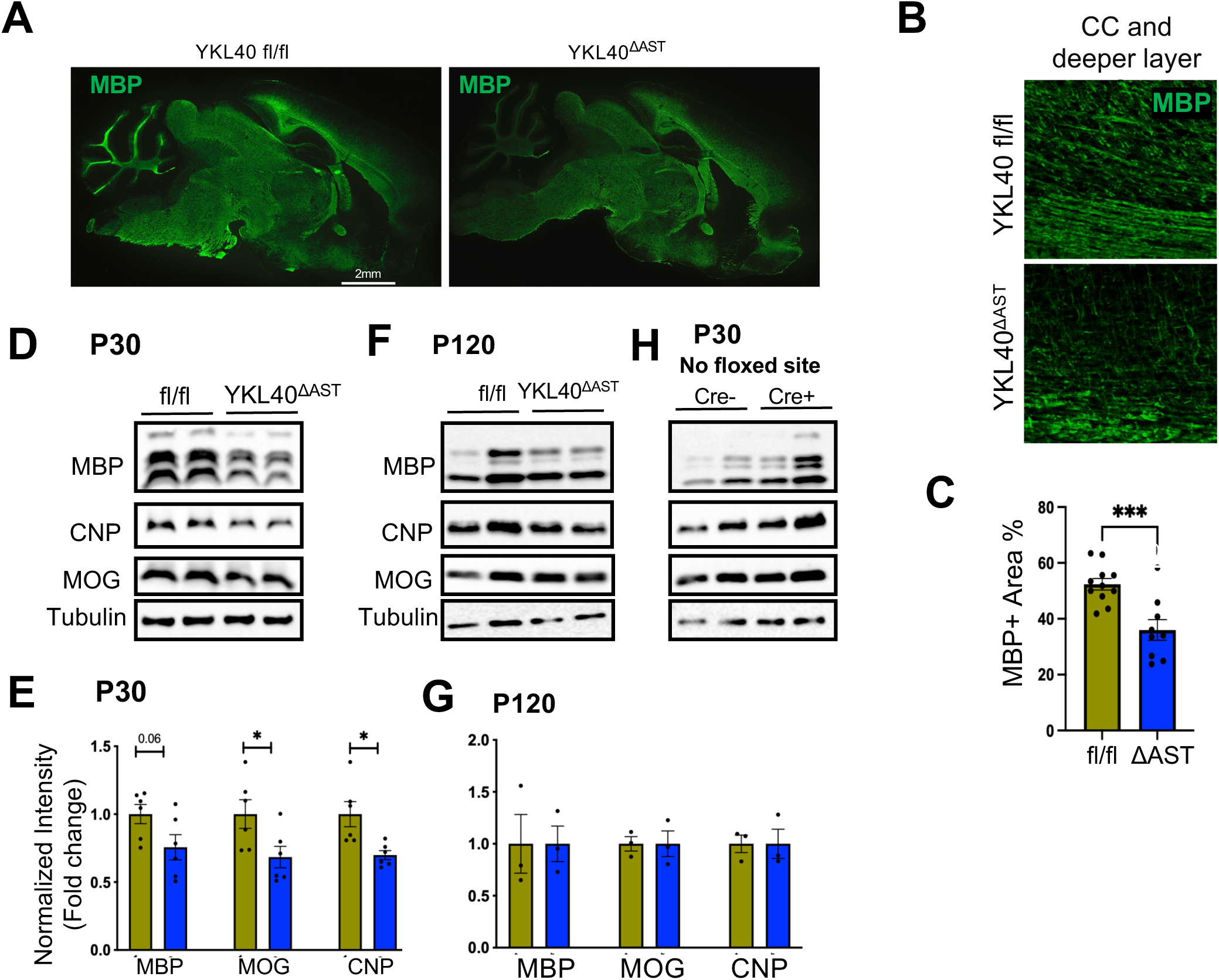
Astrocyte specific YKL40 KO mice (YKL40^ΔAST^) show delayed myelination. (**A**) Tile scan confocal image of MBP from P30 YKL40^ΔAST^ or littermates. (**B**) Representative confocal image from the CC and deeper Cx of MBP (green) and their quantification as percent area (**C**) from P30 YKL40^ΔAST^ or littermate control mice. n=10-12 images from 3,3 mice per group. (**D- G**) Western blots and its analyses of myelin markers MBP, CNP and MOG from P30 and P120 YKL40^ΔAST^ or control brains. Tubulin is used as loading control. (**H**) Western blot images of MBP, CNP and MOG from GFAP cre+/- without flox brains at P30. Tubulin is used as loading control. n=6 mice per group (E); n=3,3 (G) mice per group. Data mean ± SEM, * P < 0.05, unpaired t-test; ns=non-significant.

In the rodent brain, OL differentiation and subsequent myelination takes place postnatally starting around P6-P8, peaking at around P25-P30 and ending at around P60. We analyzed expression of mature myelinating OL marker MBP by IF in deeper cortex (CX) with corpus callosum (CC). We found a significant reduction in MBP-intensity and -percent area in this brain region (Fig. 4B,C) in YKL40^ΔAST^ compared to littermate controls. Next, we measured other well established mature OL and myelin-associated protein markers including MOG and CNPase by western blot (Fig. 4D-G). Interestingly, both MOG and CNPase protein levels were significantly reduced while MBP levels were trending towards significance in the brains of P30 YKL40^ΔAST^ (Fig. 4D,E) further confirming a role of astrocytic YKL40 in the general impact of OL maturation/myelination. Surprisingly, this myelination defect was transient since we did not observe changes in the expression of MBP, MOG or CNPase at ∼P120 YKL40^ΔAST^ brains (Fig. 4F,G), suggesting a crucial role of YKL40 in developmental myelination. To rule out any possibility of a mouse line effect on myelination we analyzed MBP, MOG and CNPase expression by WBs from the P30 brains lysates of GFAP-Cre+ or -Cre- mice without any floxed gene, and no obvious change was observed (Fig. 4H) establishing a role of astrocytic YKL40 in OL development and myelination.

Since YKL40 expression starts to increase at P8 and continues through P25 (Fig. 1A) and coincided with OL differentiation in the CC, we set out to examine OL differentiation within the CC across postnatal time P10, P16 and P30 from YKL40^ΔAST^. To do this we immunostained for CC1 a known marker of mature OLs that primarily stains the cell body, and counted cells in the CC. We found no changes in the CC1 cell counts between YKL40^ΔAST^ or littermate control mice at P10 suggesting astrocytic YKL40 does not play a role in this early OL differentiation (Fig. 5C,D). However, YKL40^ΔAST^ showed a robust decrease in CC1+ mature OLs both at P16 and at P30 compared to their littermates (Fig. 5C,D) implicating YKL40’s crucial role during peak OL differentiation. Interestingly, while CC1 positive cells robustly increased from P10 to P30 in control mice, YKL40^ΔAST^ showed lack of increase particularly between P10 and P16 (Fig. 5D). Taken together, astrocytic YKL40 is needed for OL differentiation during peak myelination.

**Fig 5.**
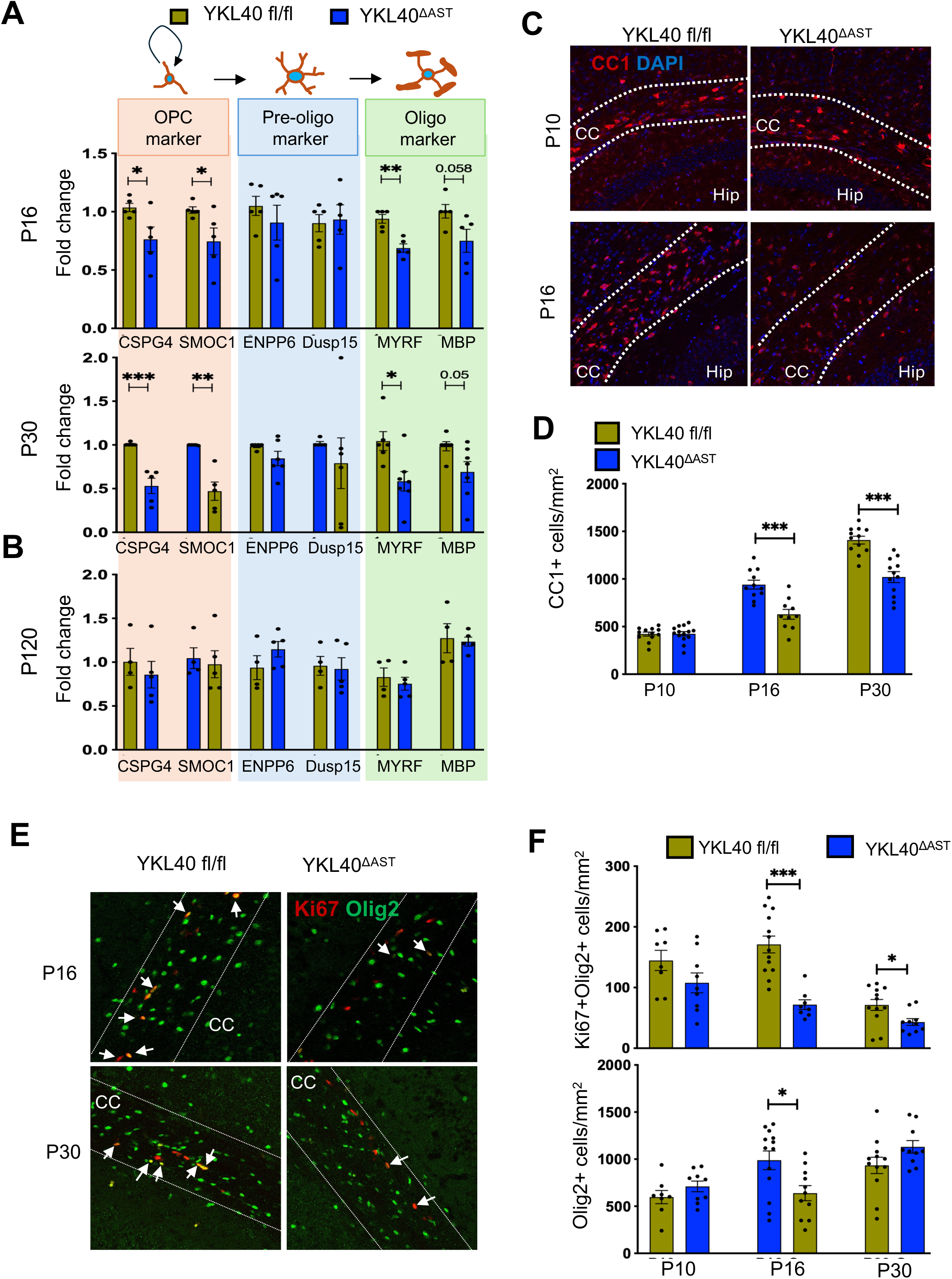
Oligodendrocyte differentiation is affected in YKL40^ΔAST^. (**A,B**) Quantitative realtime PCR analyses of expression of OL-lineage marker genes CSPG4, SMOC1, ENPP6, Dusp15, MYRF and MBP at P16 and P30 (**A**) and P120 brains (**B**). n=5,5; 6,7 (A) and n=4,5 (B) mice respectively; Data mean ± SEM, * P < 0.05, unpaired t-test. (**C,D**) Representative confocal images (**C**) and their quantification as fold change relative to GAPDH and normalized to their litter (**D**) of CC1+ maturing OLs (Red) in the corpus callossum (CC) from P10, P16 and P30 YKL40^ΔAST^ or littermate controls. (**E,F**) Representative confocal images (E) and their quantification (F) of Olig2 (Green) and Ki67 (Red) in the corpus callossum from P10, P16 and P30 YKL40^ΔAST^ or littermate controls. n=8-14 images from 3,3 mice (P10 and P16) and 3,4 mice (P30) per group. Data mean ± SEM, * P < 0.05, unpaired t-test; ns=non-significant.

### Astrocyte YKL40 is crucial for OPC proliferation and OL differentiation

Because OL-differentiation takes place in three distinct phases: i) proliferation of OPCs, ii) initial commitment of OPCs into pre-OLs and iii) final maturation of OLs from Pre-OLs and that changes in any of these stage-specific cells lead to abnormal OL differentiation, we assessed the status of OPCs, Pre-OLs and OLs in YKL40^ΔAST^ brains. To do so we analyzed expression of marker genes CSPG4 and SMOC1 (for OPCs), ENPP6 and DUSP15 (for Pre-OLs), and MYRF and MBP (for OLs) as used by others^36^. To our surprise, YKL40^ΔAST^ brains showed strong reduction in OPC markers (CSPG4 and SMOC1) in addition to reduction in mature OL markers (MYRF and MBP) at both P16 and P30 (Fig. 5A) while expression of Pre-OL marker genes remained unaltered (Fig. 5A). Similar analyses of these genes at P120 showed no changes in their levels (Fig. 5B) again suggesting hypomyelination phenotype dissipates by this age. Taken together these data indicate that YKL40 affects OL maturation by impacting OPC maintenance and OL maturation from Pre-OLs.

Reduced levels of OPCs’ marker genes in YKL40^ΔAST^ brains suggest a reduced number of OPCs which could be due to decrease in their proliferation or enhanced cell death. Since OPCs are the main proliferative cells in the CNS^37^, we first assessed OPC proliferation in the CC of YKL40^ΔAST^ mice. To examine this, we costained brain slices with pan oligo lineage marker Olig2 and a marker for dividing cells Ki67 at P10, P16 and P30 (Fig. 5E,F). We found a significant decrease in the Ki67+Olig2+ double positives cells (indicative of proliferative OPCs) at P16 and P30 in YKL40^ΔAST^ CC and minimal changes at P10 suggesting astrocytic YKL40 is needed for proper OPC proliferation. Interestingly, this decrease in OPC proliferation was more pronounced at P16 than P30 (Fig. 5E,F). In the same analyses, we found a significant decrease in overall Olig2+ cells at P16 (Fig. 5F) likely due to a decrease in populations of both OPCs and maturing OLs (Fig. 5A and F). Taken together these results strongly place astrocyte YKL40 as a crucial modulator of myelination in the developing brain by regulating OPC proliferation and inducing OL-differentiation.

### Recombinant YKL40 promotes oligodendrocyte differentiation and OPC proliferation *in vitro*

Because YKL40 is a secreted protein, the *in vivo* hypomyelination phenotype present in the YKL40^ΔAST^ brains (Fig. 3–6) may be an indirect effect on other CNS cells. To test a direct role of YKL40, we sorted to establish role of YKL40 in OL cultures *in vitro*. To do so, we first established a O4+ Pre-OL cell isolation and culture protocol from P7 mouse brains using O4 magnetic beads from Milteny biotech (Fig. 6A)^29^. This protocol yielded >90% pure Pre-OL cultures. We then applied purified YKL40 protein on these cultures for 48 hours in OL differentiation medium (Fig. 6C). Biologically relevant concentrations of purified YKL40 protein significantly increased expression of MBP, MYRF and Olig2 in a dose-dependent manner (Fig.6D). Moreover, we observed YKL40 robustly increased MBP+ cells in addition to a significant increase in Olig2 intensity in these cultures (Fig. 6E-H). Because YKL40^ΔAST^ mice displayed a reduction in OPC proliferation (Fig.5) and that O4 positive cells represent a range of proliferative cells, committed maturing cells as well as cells in between^38,39^, we also examined the effect of YKL40 on Pre-OL proliferation by using Ki67 (Fig. 6I-K). Although YKL40 treated conditions did not affect overall number of Ki67+ OPCs (Fig. 6J), we noticed reduced Ki67 intensity in many OPCs. Because cell proliferation, cell-cycle exit and cell differentiation is interrelated processes, we therefore, quantified Ki67 intensity per nuclei and found that YKL40 application significantly reduced the level of Ki67 intensity per nuclei in these cultures (Fig. 6K) indicative of more maturation state. These results again support the role of YKL40 in the maturation of pre-OLs.

**Fig 6.**
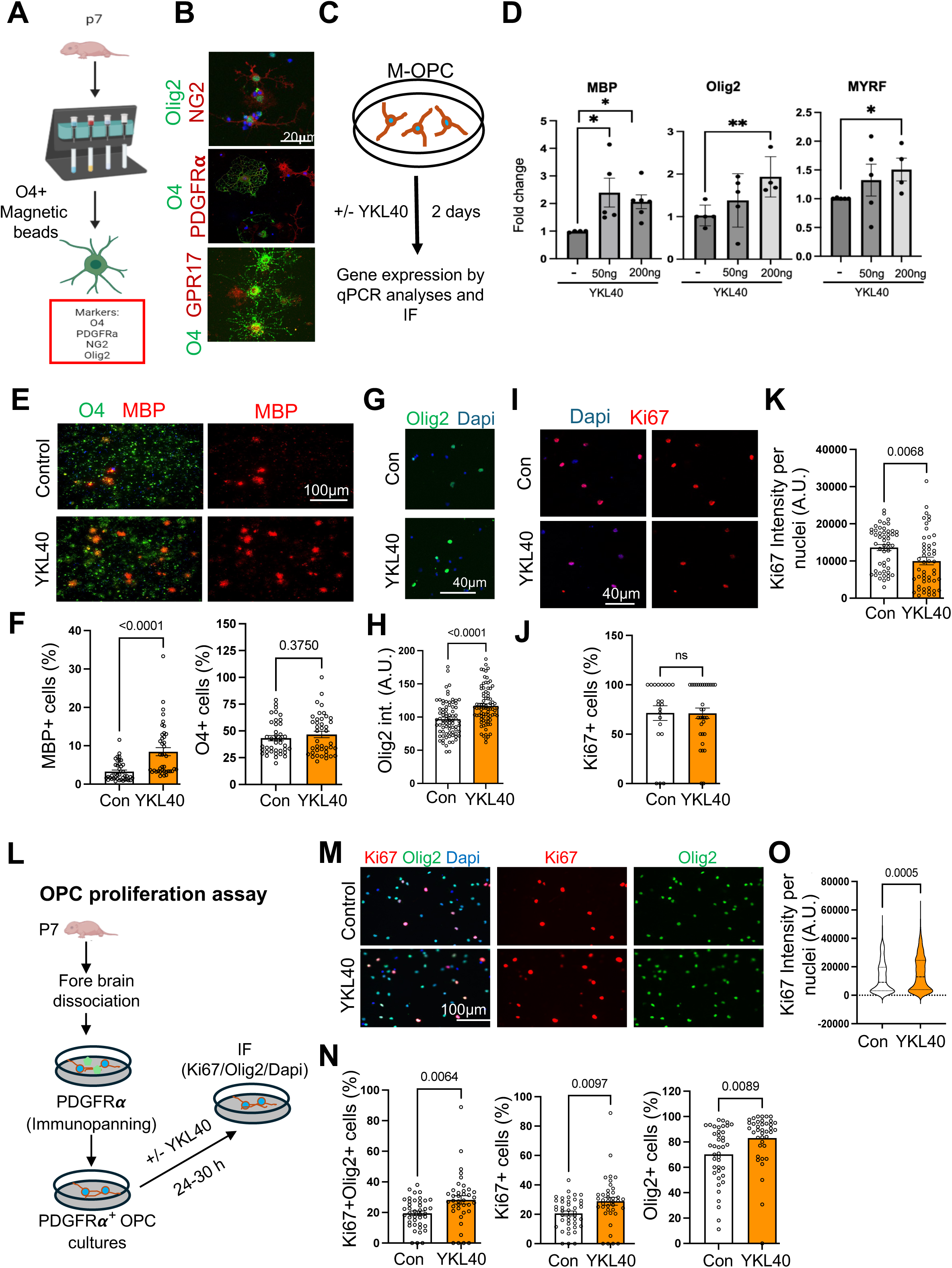
YKL40 stimulates OPC proliferation and POL differentiation. (**A,B,C**) Schematics and example of mouse O4+ POL isolation and culture with or without YKL40. (**D**) YKL40 induced expression of MBP, MYRF and Olig2 by qPCR analyses. (**C-K**) Representative confocal images of MBP, O4, Olig2 and Ki67 and their respective quantitation (F,H,J and K) of POLs incubated with or without YKL40 for two days. (**L**) Schematics and example of mouse PDGFR + OPC isolation and culture with or without YKL40. (**M,N,O**) Representative confocal images of Olig2, and Ki67 and their respective quantitation of OPCs stimulated with or without YKL40. n=>30 images from 2 biological replicates (F,H,J,K) and n=>40 images from 3 biological replicates (N,O). Data mean ± SEM, * P < 0.05, unpaired t-test; ns=non-significant.

Since, we found that YKL40 is crucial for proliferation/maintenance of OPCs *in vivo* (Fig. 5), we tested its direct role on OPC proliferation/maintenance in cultures. To do so, we isolated PDGFRα positive OPCs using immunopanning method as described in^40^ (Fig. 6L). In line with the *in vivo* data, YKL40 application increased overall number of Olig2+ Ki67+ OPCs in cultures (Fig. 6M, N). We also found an overall significant increase in Olig2+ cells by YKL40. Moreover, Ki67 intensity per nuclei also increased in OPCs by YKL40 (Fig. 6O). Taken together these data emphasize a dual role of YKL40 during OL differentiation: a) in the maintenance/proliferation of PDGFRα+ OPCs and b) the differentiation of pre-OLs into OLs.

### OPCs stimulate expression of YKL40 in astrocytes which in turn facilitates OL differentiation *in vitro*

In the developing brain, astrocytic YKL40 expression in the white matter coincides with OL development and that YKL40 is crucial for proper OL differentiation, (Fig. 1–6), we reasoned that developing OPCs may stimulate expression of YKL40 in astrocytes. To test this, we established an *in vitro* coculture system consisting of primary human astrocytes and mouse O4+ OPCs (Fig. 7A). This unique coculture system enabled us to analyze changes in gene expression in both OLs and astrocytes by using species-specific qPCR primers as we have previously reported in a human astrocyte-mouse neuronal coculture system^41^. Surprisingly, astrocytes co-cultured with O4+ OPCs for 2-days significantly induced expression of YKL40 (∼2-fold) (Fig. 7B). On the other hand, expression of other astrocyte specific genes, including GFAP, GLT1 and GLAST did not change (Fig. 7B). This suggests that OPCs do not induce expression of differentiation marker genes in astrocytes but specifically upregulate YKL40 expression. Moreover, neuronal coculture does not induce YKL40 expression in astrocytes (data not shown) highlighting OPC specific signal drives expression of YKL40 in astrocytes. On the other note, astrocytes have previously been shown to impact OPC proliferation, migration and differentiation in the developing brain^13,19,20^. We thus tested whether astrocytes influenced OPC differentiation in our coculture system (Fig. 7C). Indeed, the presence of astrocytes dramatically induced expression of both MBP, and MYRF in the OPCs compared to OPC-only cultures (Fig. 7C). These results demonstrate the presence of dynamic crosstalk between OPCs and astrocytes during developmental myelination and that astrocyte secreted YKL40 may influence OPC differentiation. To test the role of YKL40 in OPC-astrocyte crosstalk dependent myelination, we down regulated YKL40 expression in human astrocytes by SiRNA (SiYKL40) or control (SiCon) and collected their conditioned media (CM) (Fig 7D). Next, we applied this CM on OPC cultures for two days and analyzed maturation markers genes (Fig. 7D,E). Indeed, astrocyte CM devoid of YKL40 did not support OL maturation as measured by reduced levels of MBP, MYRF and Olig2 compared to SiCon CM. We validated knock down efficiency of YKL40 by qPCR (Fig. 7E). These results establish YKL40 as a central player in astrocyte-OPC crosstalk dependent OL differentiation and importance of astrocyte in developmental myelination.

**Fig 7.**
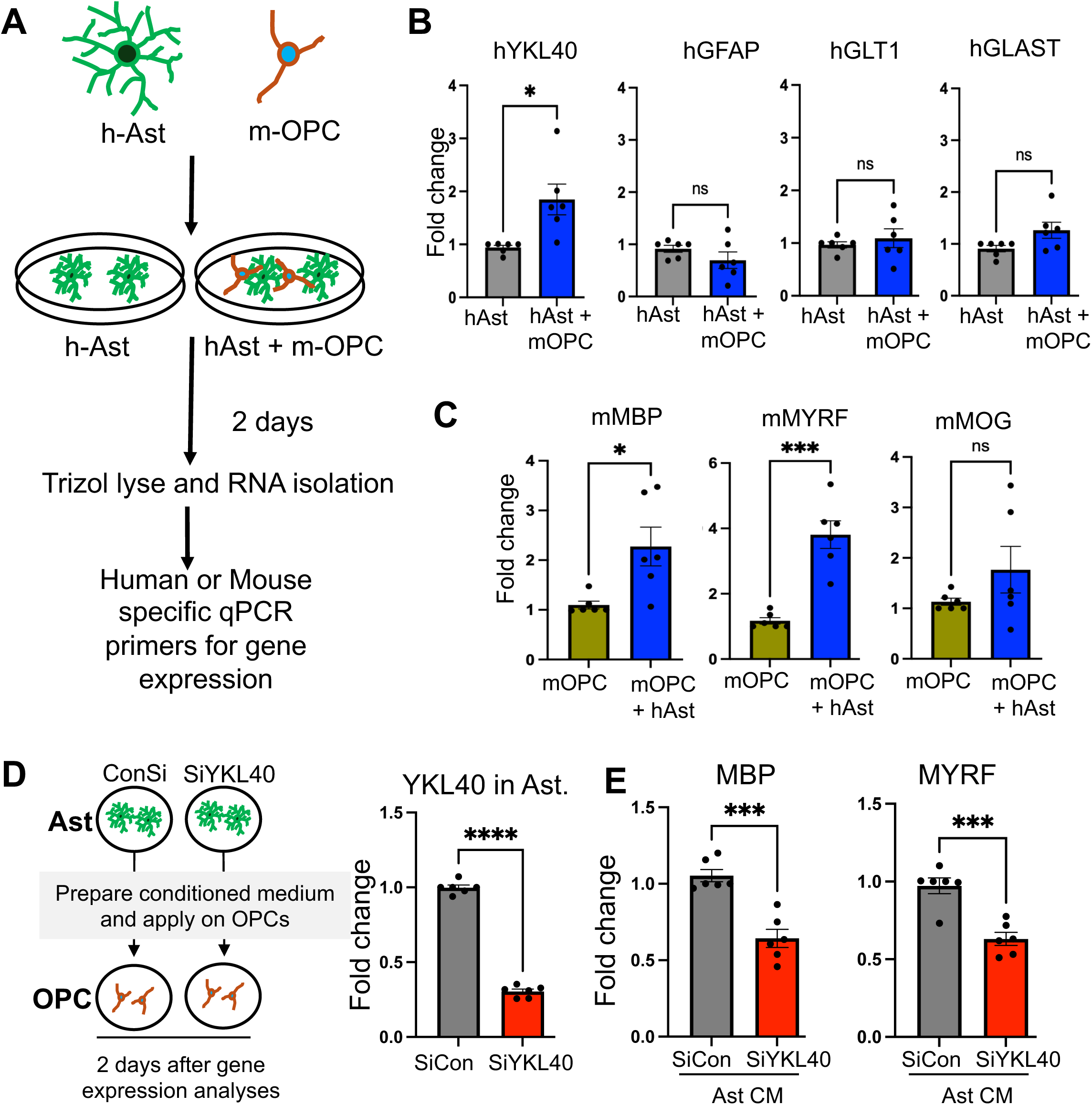
Astrocyte-OPC crosstalk induces OL-differentiation via YKL40. (**A**) Schematics of mouse O4+ POL coculture with human astrocyte to study POL-astrocyte crosstalk. (**B**) POLs induced expression of YKL40 but did not affect other astrocyte-specific genes (GFAP, GLT1 and GLAST) in astrocytes. (**C**) On the other hand, astrocytes promoted POL differentiation as measured by MBP and MYRF. n=5-6 independent cultures from 3 biological replicates. (**D**) Schematics of siRNA mediated YKL40 knock-down in astrocytes and condition media application on POL cultures. qPCR analyses confirmed YKL40 knock-down in human astrocyte cultures (right panel). (**E**) Condition media from YKL40 knock-down astrocytes does not support POL differentiation. n=6 independent cultures from 3 biological replicates. Data mean ± SEM, * P < 0.05, unpaired t-test; ns=non-significant.

## Discussion

Although YKL40 has emerged as an inflammation associated gene and correlates with disease pathology in AD, MS and glioma^23^, we do not know the fundamental role of YKL40 in the developing CNS. In this manuscript, we generated astrocyte specific YKL40 knock out mice and found that YKL40 deletion causes delayed developmental myelination by directly impacting OPC proliferation and differentiation. Astrocytes are abundant in the mammalian CNS and interact with other CNS cells to modulate brain homeostasis in health and disease. While one of the major functions of astrocytes in the CNS is to provide metabolic support to neurons and modulate synaptic function and plasticity, these cells can also interact with OLs and modulate myelination^20,42,43^. We find that O4+ magnetic bead isolated pre-OLs induce expression of YKL40 in astrocytes which in turn enhances OL differentiation. Other studies reported that YKL40 promotes OPC specification from neural stem cells^44^, while another reported to inhibit this process^45^. Similarly, one study showed that global YKL40 KO exacerbates the clinical course and pathology of EAE, whereas another showed no difference^46,47^. Moreover, intraperitoneal (i.p.) infusion of YKL40 during EAE significantly reduced pathology^48^. These contradictory results suggest YKL40 may have context dependent roles. We also find that developmental myelination defect in YKL40AST is completely normalized at the adult stage, suggesting its important role in the coordinated developmental myelination process which occurs together with neural circuit establishment and function. It is further supported by the fact that postnatal developmental myelination coincides with the expression of YKL40 (Fig.1,2) thus our data strongly supports its positive role in developmental myelination.

YKL40 is a secreted protein however mouse brain IF revealed YKL40 signal localized to major processes of astrocytes and colocalized with GFAP in the HIP and white matter astrocytes but not in cortical astrocytes (Fig. 2). We speculate that cellular location and secretion of YKL40 by astrocytes may be regulated process and depends on the brain region. In line with this YKL40 has been shown to be present in cytoplasm and nucleus of immune cells^49,50^. Absence of YKL40 in cortical astrocytic processes may indicate that cortical astrocytes either express substantially low levels of YKL40 or primarily secreted out. In future it would be important to identify role and regulation of YKL40 secretion.

Although our *in vitro* data clearly supports a direct role of YKL40 in OL differentiation and myelination, *in vivo* YKL40 dependent myelination may be dependent on other mechanisms. For example; YKL40 may influence new vascularization therefore impacting myelination^12,22,51,52^. In addition, YKL40 KO mice may display altered microglial phagocytosis which is crucial for proper developmental myelination^25,53–55^. YKL40 may also induce OL differentiation and myelination by engaging with extracellular matrix associated proteins and growth factor receptors signaling including, integrins, Syndecans, FGFR and PDGFR as has been shown for other processes^22^. This could explain YKL40’s dual role on OPC proliferation and POL- differentiation. For example, Syn-3 (syndecan-3) is highly expressed in proliferative OPCs and its expression decreases and is replaced by Syn-2 and 4 in post-mitotic differentiating POLs^56,57^. YKL40 may therefore induce OPC proliferation via Syn-3 while induced OL differentiation by interacting with Syn-2 and 4. Similarly, PI3K/Akt/mTOR pathway affects myelination by impacting OPC proliferation and OL differentiation^58^ and YKL40 is known modulator of PI3K/Akt pathway^22^. We therefore speculate that YKL40’s dual role in OL differentiation may stem from engaging with PI3K/Akt/mTOR pathway. Interestingly, astrocyte conditioned medium devoid of YKL40 does not support OL-differentiation (Fig.7) which may be due to decreased/altered levels of other astrocyte secreted signals including PDGF-AA, FGF, CNTF, LIF and BCAN in YKL40 knock down astrocyte conditioned medium^13–17^.

How does OPC induce expression of YKL40 in astrocytes? Our human astrocyte and mouse O4+OPC coculture specifically induced expression of YKL40 without impacting astrocyte maturation genes including GFAP and GLT1 (Fig7). Interestingly, Li et. al. also reported that OPCs did not induce astrocyte differentiation in cocultures with astrocytes supporting our finding^59^. Although OPCs express genes encoding secreted factors^59^ there specific role in the upregulation of astrocytic YKL40 is not known and remains to be tested. Therefore, we speculate that OPC derived signal(s) may be unique in nature to establish OPC-astrocyte crosstalk thereby inducing a specific set of genes including YKL40 needed for OPC differentiation and/or other metabolic needs. Although we do not know the exact mechanism, we speculate that it could be contact dependent signal or a soluble signal from OPCs. It will be important in the future to identify the nature of the signal and its engagement with astrocytes to induce YKL40 expression.

In conclusion, we provide the first evidence of a role of astrocytic YKL40 in developing rodent brain. Because it is highly upregulated in inflammatory conditions including MS, AD and glioma and regulates multitude of functions, it is possible that YKL40 may modulate several biological processes including myelination, neuronal death, and reactive gliosis depending on the context. Therefore, our current studies defining the role of YKL40 in astrocyte-oligodendrocyte crosstalk dependent myelination may have important implications on normal brain development and in brain disorders.

## Acknowledgements

This work was supported in part by NIH R01NS126504 to S.K.S and VCU-School of Medicine Startup funds to S.K.S and Massey Cancer Center support grant P3- CA106059. Microscopy was performed at the VCU microscopy Facility, supported, in part, by funding from NIH-NCI Cancer Center Support Grant P30 CA016059.

## Author Contributions

SKS, OD, and SM designed the experiments. OD performed most of the experiments with help from SM, SCRT, CT, JPG, and SKS. OD, SM and SKS analyzed data and arranged the manuscript figures. TK, BF and UO provided critical reagents. SKS wrote the manuscript with help from OD. All authors reviewed the manuscript.

## Competing interests

The authors declare that they have no competing interests.

## Availability of data and material

The data that support the findings of this study are available from the corresponding author upon reasonable request.

## Ethics approval

Mice were housed at Virginia Commonwealth University according to guidelines of the Institutional Animal Care Use Committee (IACUC). The mouse protocols were approved by the IACUC. All mice were housed with food and water available ad libitum under a 12 h–12 h light– dark cycle in a 20–22°C and 40–60% humidity environment.

**Figure Supplement 1: YKL40 is specifically expressed in astrocytes.** (**A**) YKL40 (*Chil1*) mRNA is highly expressed in purified astrocytes at P30 compared to other CNS cells in the mouse. Average FPKM of Chil31 mRNA from brainrnaseq.org. (**B**) YKL40 (CHI3L1) mRNA is specifically expressed in mature astrocytes isolated from human brain. Average FPKM of Chi3l1 mRNA from Zhang. Et. al. 2016 and brainrnaseq.org. (**C**) Chi3l1 mRNA is actively translated. Average FPKM values of Chi3l1 mRNA engaged within translating astrocyte-specific and all ribosomes (input) from Doyle et. al. 2008. (**D**) Chi3l1 mRNA is actively translated. Average FPKM values of Chi3l1 mRNA engaged within translating astrocyte-specific and all ribosomes (input) from Srinivasan et. al. 2016. (**E**) YKL40 mRNA visualized by FISH (Red) in hippocampus and corpus callosum astrocytes (green) from P30 ALDH1L1.eGFP reporter mouse. 3-D Imaris rendering and surfaces of an individual astrocyte and YKL40 mRNA (red dot) FISH signal (lower panel).

## Notes

### Competing Interest Statement

The authors have declared no competing interest.

