## Supplementary figures and images for "Astrocyte-oligodendrocyte crosstalk dependent myelination via secreted protein YKL40"

### figure supplement S1

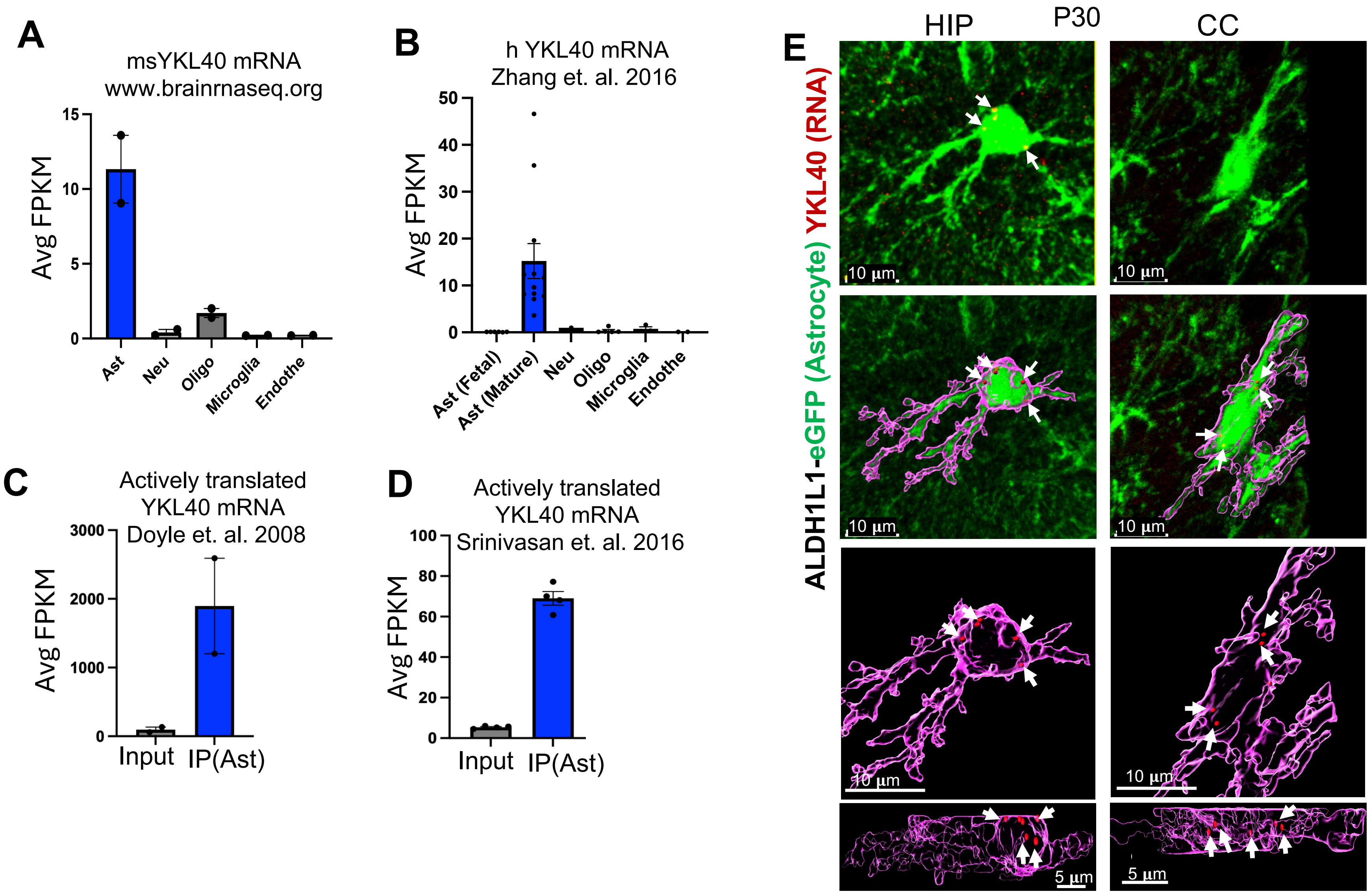
